# Prioritization of regulatory variants with tissue-specific function in the non-coding regions of human genome

**DOI:** 10.1101/2021.03.09.434619

**Authors:** Shengcheng Dong, Alan P. Boyle

## Abstract

Understanding the functional consequences of genetic variation in the non-coding regions of the human genome remains a challenge. We introduce here a computational tool, TURF, to prioritize regulatory variants with tissue-specific function by leveraging evidence from functional genomics experiments, including over three thousand functional genomics datasets from the ENCODE project provided in the RegulomeDB database. TURF is able to generate prediction scores at both organism and tissue/organ-specific levels for any non-coding variant on the genome. We present that TURF has an overall top performance in prediction by using validated variants from MPRA experiments. We also demonstrate how TURF can pick out the regulatory variants with tissue-specific function over a candidate list from associate studies. Furthermore, we found that various GWAS traits showed the enrichment of regulatory variants predicted by TURF scores in the trait-relevant organs, which indicates that these variants can be a valuable source for future studies.

## Introduction

Characterizing the biological impact of variation in the non-coding regions of the human genome remains a challenge in the interpretation of human diversity. Genome-wide association studies (GWAS) have identified millions of genetic variants that are associated with diverse disease traits (Buniello et al. 2019). Most of these variants (∼90%) map to the non-coding regions of the human genome (Tam et al. 2019). Due to the lack of understanding of these regulatory elements within non-coding regions, it is important to assess the functional consequences of these disease-related variants from GWAS.

To facilitate studies of non-coding genomic regions, large consortia, including ENCODE (Davis et al. 2018; ENCODE Project Consortium 2012) and the Roadmap Epigenomics projects, (Bernstein et al. 2010) have defined the human regulatory landscape using high-throughput functional genomics assays. For example, DNase-seq locates open chromatin regions in the genome (Boyle et al. 2008; Song and Crawford 2010), while ChIP-seq identifies chromatin modification patterns and transcription factor (TF) binding sites within regulatory elements (Barski et al. 2007; Johnson et al. 2007; Mikkelsen et al. 2007). With further incorporation of variant genotypes into these methods, variants associated with differential TF binding and chromatin states have been described (Kasowski et al. 2013, 2010; Degner et al. 2012). In addition, massively parallel reporter assays (MPRA) identify regulatory variants that affect gene expression levels directly (Tewhey et al. 2018; Kheradpour et al. 2013; Patwardhan et al. 2012). These studies demonstrate that a significant number of variants drive regulatory state variation across the population, and potentially explain the diversity in disease risk and phenotype observed from GWAS studies.

Computational tools have helped prioritize regulatory variants in non-coding regions by leveraging knowledge from functional genomics assays. Prediction scores of functional probability for variants are available from tools including RegulomeDB (Boyle et al. 2012), GWAS3D (Li et al. 2013), HaploReg (Ward and Kellis 2016), DeepSEA (Zhou and Troyanskaya 2015), DeepBind (Alipanahi et al. 2015), DanQ (Quang and Xie 2016) and Basenji (Kelley et al. 2018). The process of narrowing down a candidate list of variants using these prediction scores can reduce time-consuming validation experiments. However, most current computational tools overlook the uniqueness of gene regulatory networks found within different tissues by only providing a prediction score at an organism level. This can be misleading for research groups focused on tissue-specific functional variants. New tools have recently become available that provide tissue-specific prediction scores, such as FUN-LAD (Backenroth et al. 2018), GenoNet (He et al. 2018), cepip (Li et al. 2017) and GenoSkyline (Lu et al. 2016). However, they mainly utilize epigenetic data from the Roadmap Epigenomics project (Bernstein et al. 2010) making it hard to leverage their results against other tissues not included in the Roadmap project. The ENCODE project currently houses thousands of ChIP-seq and DNase-seq datasets in over 200 tissues and cell types, including those from the Roadmap project, that can further increase the scale and accuracy of tissue-specific function prediction.

Here we introduce a computational tool, TURF (Tissue-specific Unified Regulatory Features), that prioritizes regulatory variants in the non-coding regions of the human genome. TURF is built on our RegulomeDB framework to allow for easy delivery of our predictions as well as constant updates in the functional annotations across the human genome. We extend our previous algorithm SURF (Dong and Boyle 2019) to predict tissue-specific functional variants in addition to the tool’s original generic context at an organism level. To construct a high-quality training set, we called 7,530 allele-specific TF binding (ASB) single nucleotide variants (SNVs) in 6 cell lines from over 600 ChIP-seq datasets. We then trained a random forest model using features from functional genomic annotations across all available tissues from ENCODE. This classifier greatly improves the robustness of RegulomeDB v1.1 ranking scores and surpasses other top-performing tools on an independent MPRA dataset. We then incorporated annotations of histone marks and open chromatin regions in a particular tissue to train a separate random forest model and obtain a final tissue-specific score. The tissue-specific score leverages information from other tissues, as well as retaining the uniqueness of individual tissues. Moreover, we extended the tissue-specific scores to organ-specific scores in the 51 organs with available genomics data from the ENCODE project. The pre-calculated organ-specific scores for all GWAS SNVs from the GWAS Catalog are available at https://github.com/Boyle-Lab/RegulomeDB-TURF and TURF is currently being integrated into RegulomeDB v2.0.

## Results

### Overview of the TURF algorithm

TURF prioritizes non-coding variants with both generic scores and tissue-specific scores (Fig. 1). It first uses a random forest model built by training on features from functional genomics annotations in all available tissues and cell types from the ENCODE project (Davis et al. 2018). It uses a similar feature set to our previously successful algorithm SURF (Dong and Boyle 2019), including binary features retrieved from the original RegulomeDB ranking scheme and functional significance scores from DeepSEA (Zhou and Troyanskaya 2015). Furthermore, it includes continuous signals from ChIP-seq assays to increase the resolution of the algorithm (see features list in Supplemental Table S1). Generic scores from the first random forest model predict whether the query variant is functional in any human tissue. Tissue-specificity is further predicted by using a separate random forest model trained on functional genomic annotation features only from a particular tissue. To avoid data availability bias for different tissues, TURF takes advantage of DNase-seq and well-studied histone mark ChIP-seq data that cover most tissues (see features list in Supplemental Table S1). By combining the probability score from the second random forest model with the generic score from the first model, the resulting tissue-specific score predicts the probability of the query variant being functional in a specific tissue.

**Figure 1.**
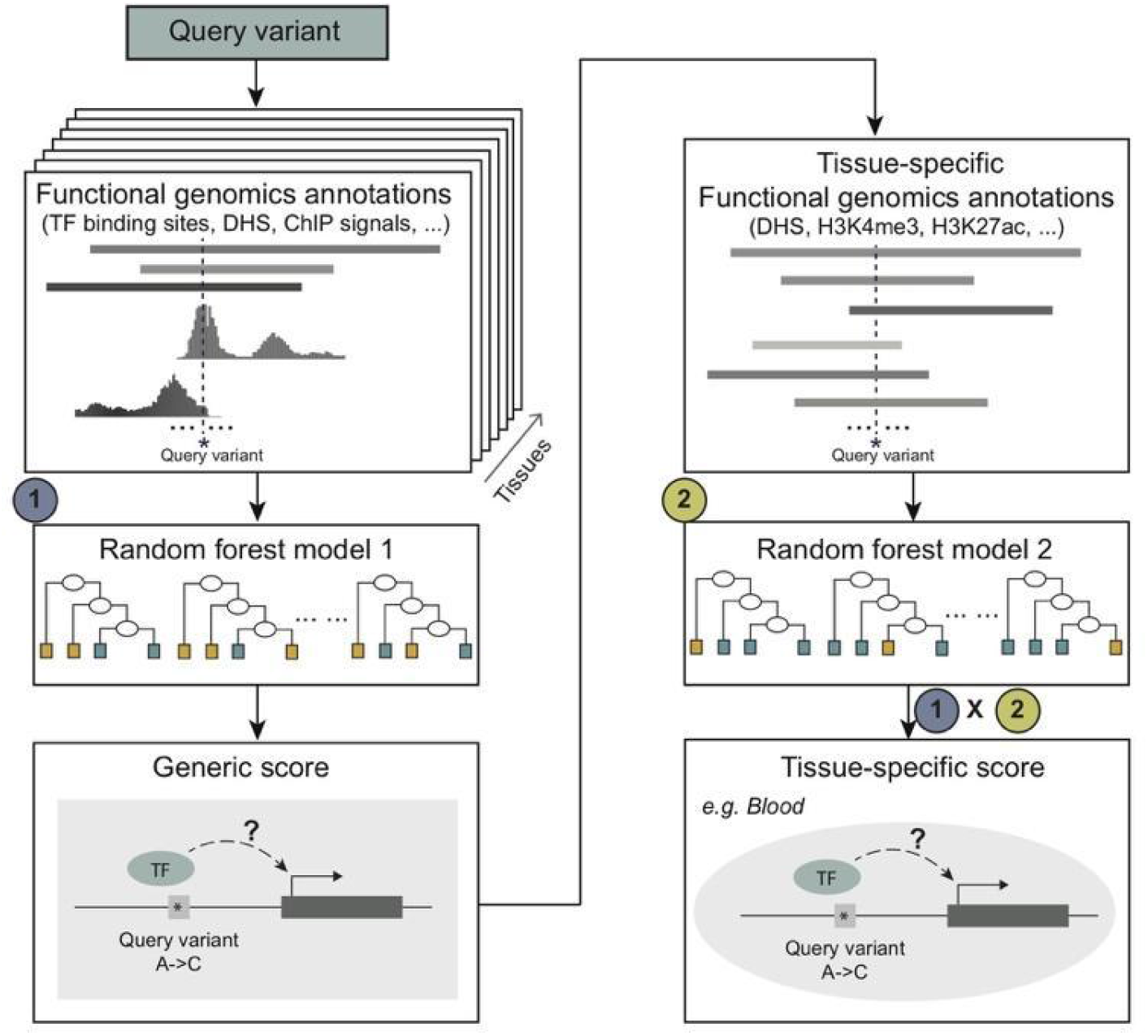
Overview of TURF algorithm. TURF generic score predicts the probability of a query variant being functional in any tissue from the first random forest, which used features of functional genomics annotations from all available tissues. By further incorporating annotations from a given tissue, a tissue-specific prediction score is computed by multiplying generic score with the prediction score from a second random forest model.

### TURF generic score improves the performance of RegulomeDB v1.1 ranking score

TURF improves on the original heuristic ranking score in RegulomeDB v1.1 by providing a probabilistic score generated from a random forest model. By replacing the single empirical decision with sets of decision trees, the model avoids issues caused by excessive reliance on only a few functional genomic annotations. To develop a training set for the model, we generated a set of variants with high confidence functional confidence through identification of 7,530 allele specific transcription factor binding (ASB) single nucleotide variants (SNVs) in six cell lines (GM12878, HepG2, A549, K562, MCF7 and H1hESC) by reprocessing 864 ChIP-seq datasets from the ENCODE project using *AlleleDB* v2.0 (Chen et al. 2016). ASB SNVs were called if different TF binding affinity with a single nucleotide change at heterozygous sites was observed. We defined a background set using non-allele specific TF binding SNVs as well as a set of variants outside TF binding regions (see methods).

We evaluated the TURF generic score performance on an independent and orthogonal dataset from a massively parallel reporter assay (MPRA) (Tewhey et al. 2018). This dataset was also utilized as a test set in a previous paper (Zhang et al. 2019), where the authors found DeepSEA scores provided the best prediction model for calling variants functional in tissues. TURF outperformed DeepSEA scores on this MPRA test set with a larger AUROC and the same AUPR (Fig. 2). To compare with the original ranking score from RegulomeDB v1.1, we calculated TURF generic scores for all common SNPs from dbSNP153 (Sherry 2001). The SNPs that originally scored in the highest category, which was largely dominated by eQTL evidence, now show a wider range of scores that better predicted their functionalities, while the overall trend was unchanged (Supplemental Fig. S1).

**Figure 2.**
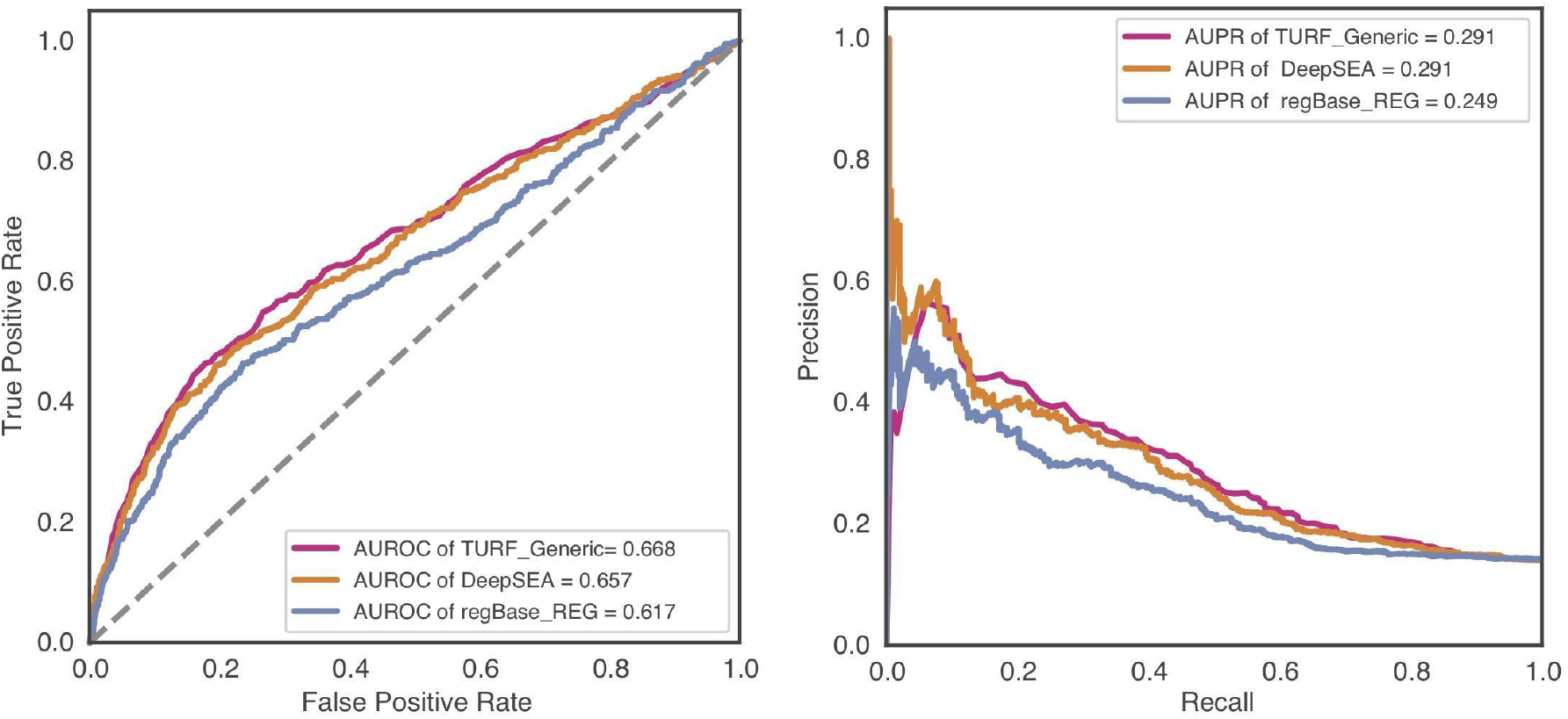
TURF generic scores performance on test data from massively parallel reporter assay (MPRA) in GM12878. Performance was evaluated by Area Under ROC Curve (AUROC) and Area Under Precision-Recall Curve (AUPR). 435 positive variants vs 2670 control variants were called in this MPRA validated dataset.

### TURF tissue-specific scores performance on MPRA data in three cell lines

We further evaluated TURF tissue-specific predictions with MPRA datasets from three cell lines (GM12878, HepG2 and K562) using the same strategy as He et al. (He et al. 2018). Tissue-specific predictions by TURF had the best performance in GM12878 versus other top performing computational tools (Fig. 3A and Supplemental Table S3). TURF also has the top AUROC in HepG2 with the second largest AUPR (0.571 compared to 0.572 from GenoNet) and the largest AUPR in K562. Noticeably, the tissue-specific features in the second random forest model have significantly improved the performance of the TURF generic scores. Among all tissue-specific features, open chromatin regions from DNase-seq in the corresponding cell lines are the most important predictors in all three MPRA datasets. Tissue-specific DNase footprints and active histone marks, including H3K4me2, H3K4me3 and H3K27ac, also play essential roles in variant prediction (Supplemental Fig. S2).

**Figure 3.**
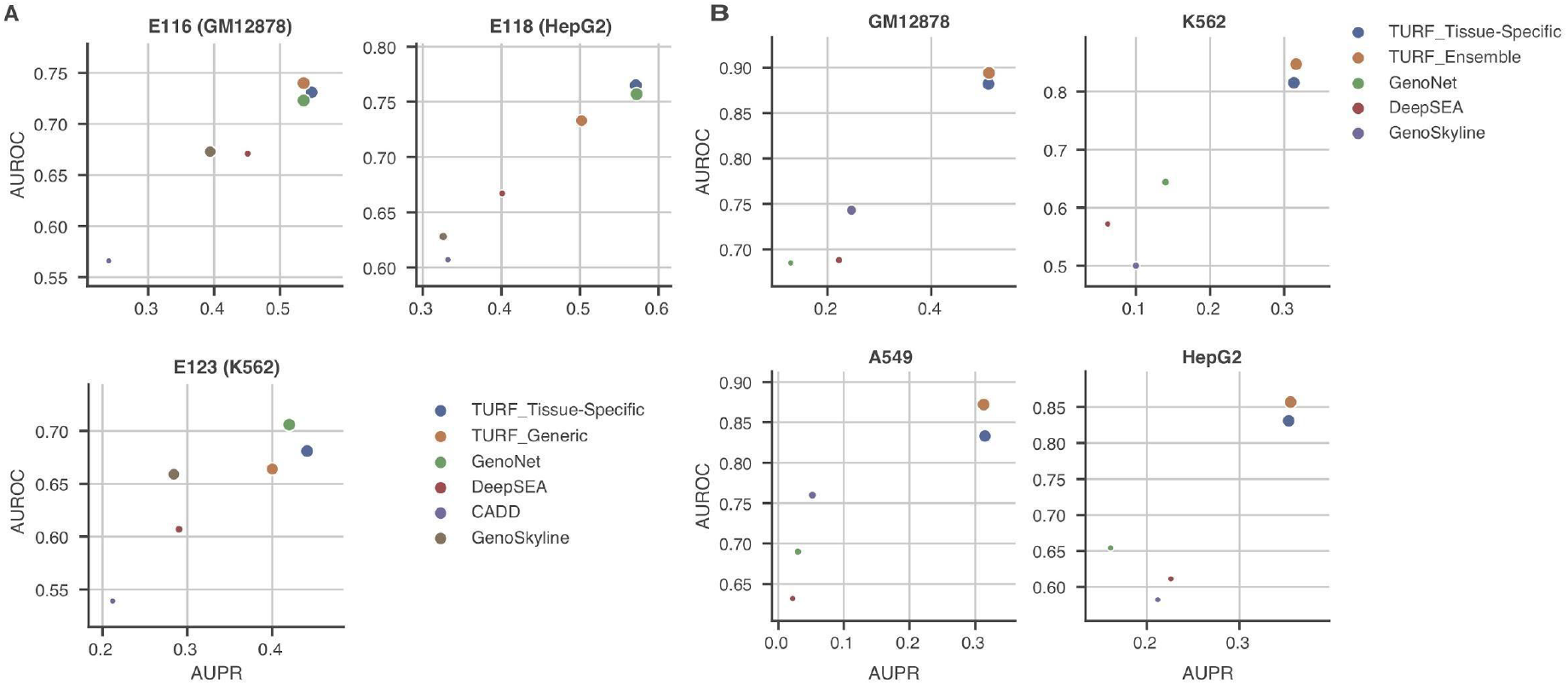
Tissue-specific predictions performance comparisons. Each plot shows the AUPR (area under the precision recall curve) on x axis and the AUROC (area under the receiver operating characteristics curve) on y axis. The size of each point represents the pearson correlation. (A) Performance on MPRA data in three cell lines (GM12878: 693 positive variants, 2772 control variants; HepG2: 524 positive variants, 1439 control variants: K562: 339 positive variants, 1361 control variants). (B) Performance on allele specific transcription factor binding SNVs (see the number of variants in Supplemental Table S2).

### TURF tissue-specific predictions on allele specific TF binding (ASB) SNVs

Despite the power of using MPRA datasets as training sets, they are currently limited in terms of the number of tested variants and the variety of tissues. To obtain a more robust tissue-specific model, we called allele-specific TF binding (ASB) SNVs from 6 cell lines. When trained on ASB SNVS, our tissue-specific models greatly outperformed other methods (Fig. 3B and Supplemental Table S3). Among the tissue-specific features, DNase-seq peaks and several active histone marks, such as H3K4me2 and H3K27ac, were important predictors of tissue-specific functional variants, similar to what was observed in the MPRA datasets (Supplemental Fig. S3). However, DNase footprints show more variation in feature importance ranking within the 6 cell lines. This indicates the diversity of DNase-seq data quality in different cell lines, and suggests that utilization of a more robust model to compensate for this variation is needed when extending to other tissues not used in the training data.

We then trained an ensemble tissue-specific model using the average predictions from 6 models with feature weights individually learnt from 6 ASB cell lines. The histone mark features were restricted to 5 histone marks that ranked high in feature importance, and had available datasets covering most tissues (i.e. H3K27ac, H3K36me3, H3K4me1, H3K4me3 and H3K27me3). The ensemble model outperformed the individual tissue-specific models when predicting ASB SNVs (Fig. 3B and Supplemental Table S3). Moreover, this ensemble model trained on ASB SNVs performed better than most of the other tools when tested on the previous independent MPRA datasets in all three cell lines. The exception was GenoNet, which used labels from the MPRA datasets in their training step (Supplemental Table S3). Predictions were computed from this ensemble tissue-specific model on the ASB SNVs in 6 cell types and most exhibited the highest prediction scores in their corresponding functional cell line (Fig. 4). However, HepG2 ASB SNVs had the least enrichment of high HepG2-specific scores, perhaps due to DNase-seq noise in the dataset as only 25% were in DNase peaks. Some H1hESC ASB SNVs had high scores in K562 and MCF7, implying that a many stem cell regulatory variants are involved in regulation of pathways in differentiated cell lines.

**Figure 4.**
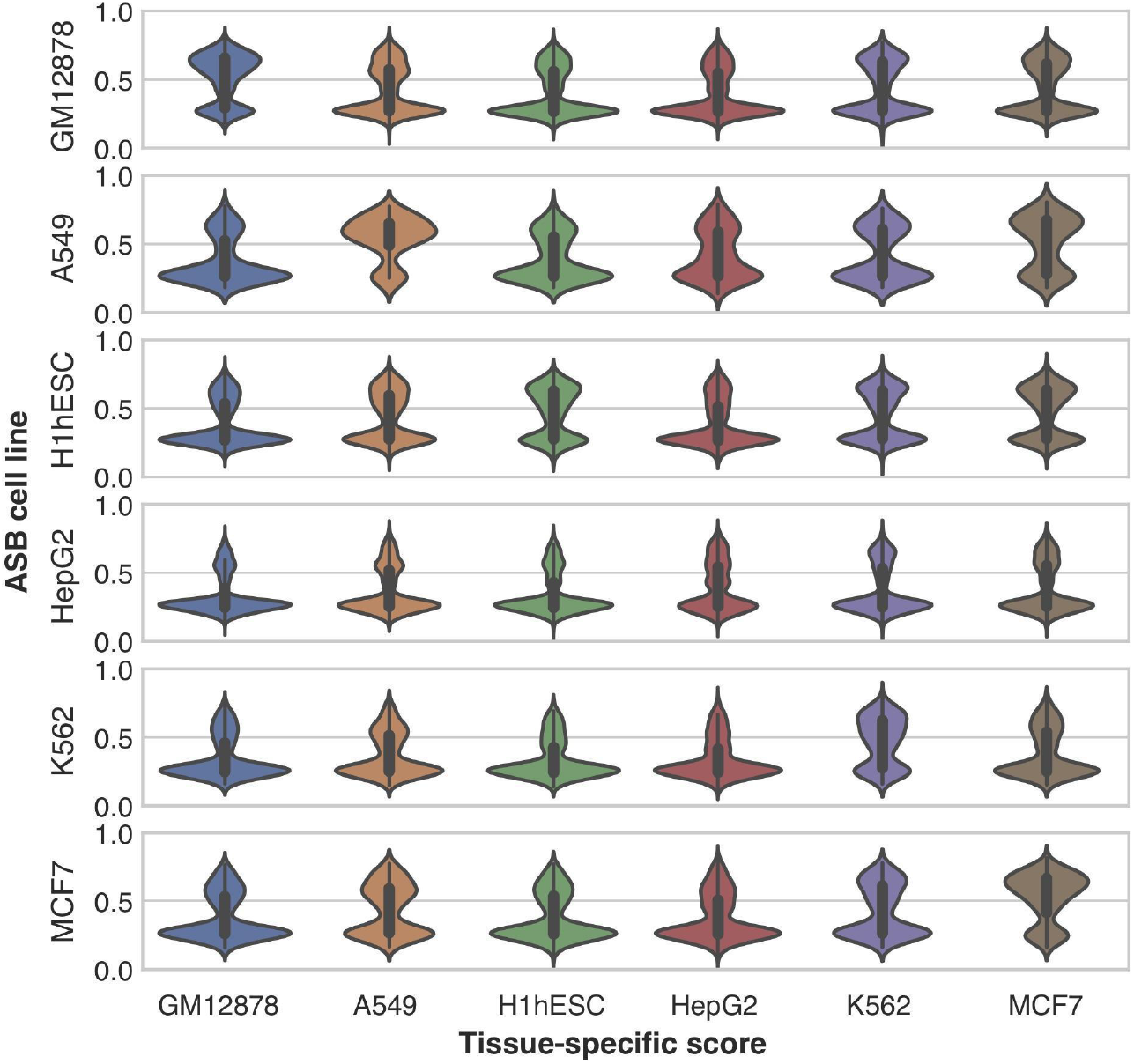
TURF tissue-specific scores on allele-specific transcription factor binding (ASB) SNVs called from 6 cell lines. The ASB cell line represents the functional tissue for ASB SNVs in each row. The tissue-specific scores are shown in violin plots with a given cell line in each column. ASB SNVs have overall the highest tissue-specific scores in their functional cell line.

### Extension of TURF tissue-specific scores to organ-specific scores

To expand the scale of prediction for TURF, we leveraged tissue-specific functional genomic annotations of tissues belonging to the same organ and generated combined organ-specific scores across 51 organs. We were able to recover the organ-specific function of some well-studied regulatory variants in specific genomic loci with TURF scores. For example, TURF’s organ-specific scoring was able to pick out the regulatory SNP rs12740374 that affects liver-specific *SORT1* gene expression levels in the 1p13 cholesterol locus (Musunuru et al. 2010) (Fig. 5). The liver-specific function of rs12740374 was also validated in HepG2 MPRA assays (Shigaki et al. 2019). The position of rs12740374 overlaps several active histone mark peaks from ChIP-seq (H3K27ac, H3K4me3 and H3K4me1) and DNase peaks in liver tissues. These multiple lines of genomics evidence prioritized rs12740374 as the top SNP for liver-specific scores within a list of candidates from previous association studies. In addition to liver, rs12740374 has a high significance score in other organs relevant with cholesterol metabolism, such as adipose tissue and gonad. As another example, TURF also detected a regulatory SNP at the *GATA4* locus in the heart (Supplemental Fig. S4) that was initially discovered in a genome-wide association scan on 466 bicuspid aortic valve cases (Yang et al. 2017).

**Figure 5.**
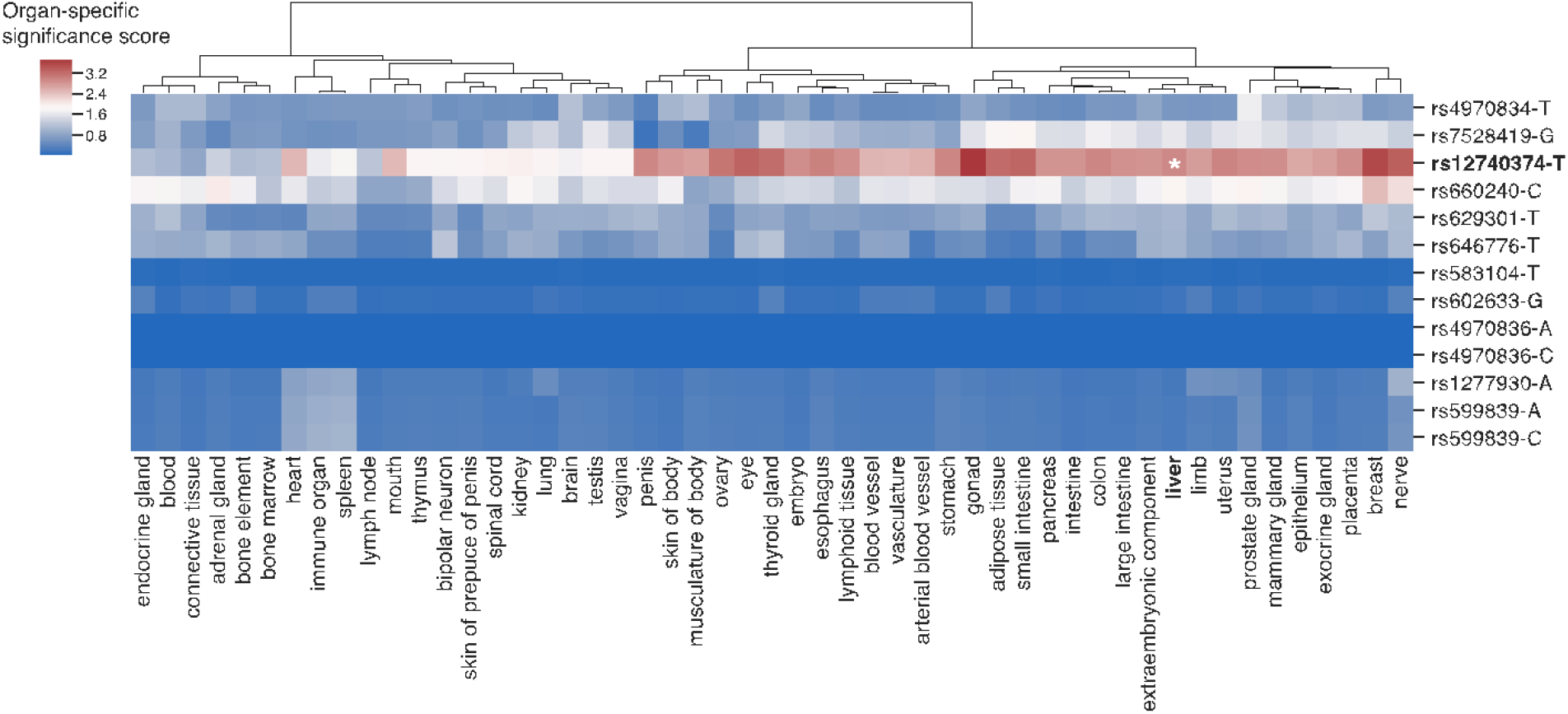
Organ-specific significance scores of variants in the 1p13 cholesterol locus. rs12740374 has the top liver-specific significance score compared to other nearby candidate SNPs from association studies, which was validated to affect *SORT1* gene expression level in liver tissue. The organ-specific significance scores were calculated relative to a background set from GWAS variants (see methods).

### TURF organ-specific scores prioritize genetic variants associated with traits in relevant organs

We examined TURF organ-specific scores on variants identified from genome-wide association studies (GWAS) using the GWAS Catalog portal (Buniello et al. 2019). GWAS variants were found to be enriched in regulatory elements of non-coding regions (Maurano et al. 2012; Roadmap Epigenomics Consortium et al. 2015). We tested the enrichment of putative regulatory variants prioritized by TURF scores for a variety of traits. For each trait, the top organ with the highest z-score showed the most significant enrichment of organ-specific regulatory variants relative to the background set from all traits within the GWAS catalog, as well as 50 other organs with the same trait (Fig. 6 and Supplemental Fig. S5).

The top enriched organs from diverse traits were consistent with current trait-relevant organ knowledge. For example, many immune system related diseases, such as autoimmune disease, celiac disease and chronic lymphocytic leukemia, showed a high enrichment for regulatory variants functional in immune-related organs, including immune organ, spleen, and lymph node. Traits of immune cells, such as leukocyte, eosinophil and platelet, were also enriched in immune organs. Cardiac traits, including PR interval, which is a measurement in electrocardiography, and coronary artery disease, were enriched in heart and arterial blood vessel. Enrichment in the colon and immune-related organs was demonstrated for Crohn’s disease and ulcerative colitis, both inflammatory bowel diseases. Furthermore, several traits of measurement were enriched for organs involved in relevant metabolic pathways, such as cholesterol measurement in liver, apolipoprotein A1 measurement in small intestine (Glickman and Green 1977), renin-angiotensin system (RAS) use measurement in adrenal gland, and alcohol consumption measurement in exocrine gland (i.e. salivary gland). Of note, the enrichment of variants in some traits could be affected by cofactors, such as gender for body height enrichment within the vagina and ovary. Also, some organs seem to share similarities in gene regulatory networks, partly due to overlapping of tissues, or tissues with similar functions across different organs. This explains a mixture of brain and optic traits enriched in either brain or eye, as the optic nerve gene expression pattern was found to be similar to brain tissue (Diehn et al. 2005).

The most enriched organ for potential regulatory variants provides new directions for understudied diseases or traits. For instance, drugs of calcium channel blockers were found to increase the risk of pancreatic cancer in post-menopausal women (Wang et al. 2018), while the underlying mechanisms remain unclear. Interestingly, pancreas was the top organ for the calcium channel blocker use measurement trait, which indicates an enrichment of putative regulatory variants functional in pancreas. Thus, additional studies on top variants prioritized by TURF pancreas-specific scores may help further explain the association between pancreatic cancer risk and the use of calcium channel blocker drugs. Similar workflow can be applied to other diseases, such as Alzheimer’s disease in immune organs, to determine the causal variants in non-coding regions.

## Discussion

In this study, we developed TURF, a computational tool that prioritizes variants in non-coding regions. Evidence was incorporated from various functional genomic assays to produce robust predictions that were verified via MPRA assays in both generic and tissue-specific contexts. The workflow was designed to identify regulatory variants from association studies with tissue/organ-specific regulatory function. Moreover, we found GWAS variants were enriched with regulatory variants predicted by TURF organ-specific scores in trait-related organs.

To balance between prediction accuracy and data availability, we trained TURF on ASB SNVs identified from ChIP-seq to determine the weight of features in a tissue-specific context, then extended the scale of annotation to an organ-specific level. The TURF tissue-specific scores leverage information gained from other tissues while retaining the uniqueness of the gene regulatory network in individual tissues. We were able to prioritize putative organ-specific regulatory variants across 51 organs in diverse pathways. A number of computational tools have been developed recently for similar purposes however, most focus on genomic assays and tissues from the Roadmap project (He et al. 2018; Lu et al. 2016). This makes it difficult to utilize their results for tissues not included in the Roadmap project. As an alternative, we took advantage of over 3,000 genomic assays in more than 200 tissues and cell types available from the ENCODE project, expanding the annotation scope and enhancing the robustness of our predictions. Most relevant organs of various GWAS traits were recovered from the organ-specific scores, including some well-studied traits, such as LDL cholesterol measurement and immune diseases. These results were mirrored in active histone marks using epigenomics data from the Roadmap project (Roadmap Epigenomics Consortium et al. 2015). In addition, we observed novel organ-trait pairs, including pancreas in calcium channel blocker use measurement, which can help elucidate underlying disease mechanisms. As more functional genomics datasets are generated, our algorithm is flexible allowing for addition of new tissues by querying histone mark and DNase features within the new tissue and then computing new tissue/organ-specific scores.

Despite the large scale of annotation utilizing the 51 ENCODE organs, further refinement of the organ terms and the tissues assigned to each organ is possible. Some traits in Figure 5 showed enrichment in non-relevant organs, such as household income in the bipolar neuron. This could be partly due to cofactors within individual GWAS samples, but can also imply an imbalance in the number of genomic datasets across diverse organs as the bipolar neuron (i.e. ear) only contains one ENCODE biosample. Due to the limitation of data availability, we only used 7 tissue-specific binary features when building the second random forest model. With more functional genomics data being generated, especially those targeting more histone marks, we can expand our feature set and generate a wider spectrum of prediction scores. The organ-specific scores can then be normalized across different organs to eliminate bias from data availability. The organ-specific scores for a variant will be more comparable over a list of interested organs.

We used MPRA data to validate our method as these assays provide more direct evidence of variants affecting gene expression than other association analyses, such as eQTLs, which can be affected by variants that are in strong linkage equilibrium. However, we could only test our model in three MPRA cell lines when comparing performance to other tools. We found one tool used MPRA data labels causing overfitting when tested on ASB SNVs. We built an ensemble model trained on SNVs from 6 cell lines to avoid the overfitting. With more MPRA data becoming available in the future, we can provide a more thorough comparison of performance and further refine our model by including training variants from more cell types or more types of assays.

Overall, TURF is able to prioritize regulatory variants with either generic or tissue-specific functions. We expect our tool to enhance future studies on functional consequences of regulatory variants associated with diseases from GWAS. The organ-specific scores generated here will be incorporated into the RegulomeDB database soon, making it a useful tool for broad communities.

## Methods

### Training dataset generation

We identified 7,530 allele specific transcription factor (TF) binding (ASB) SNVs in 6 cell lines (GM12878, HepG2, A549, K562, MCF7 and H1hESC), which are defined as variants that result in stronger binding of a TF to one allele at heterozygous sites in an individual. The *AlleleDB* (Chen et al., 2016) protocol was used to call ASB SNVs.

The SNVs in GM12878 and H1hESC were obtained from the 1000 Genome Project (1000 Genomes Project Consortium et al. 2015) and NCBI GEO database (accession number: GSE52457) separately. For the other four cell lines, variants were called from their whole genome sequencing data (data accessible at NCBI SRA database with accession numbers: DRX015191, SRX2598759, SRX285595 and SRX1705314) by *HaplotypeCaller* from the Genome Analysis Toolkit (GATK) v3.6 (DePristo et al. 2011) following GATK’s Best Practices (https://gatk.broadinstitute.org/). Their diploid personal genomes were constructed using *vcf2diploid* v0.2.6 (Rozowsky et al. 2011) to avoid alignment biases favoring reads containing reference alleles by mapping to maternal and paternal genomes separately. Copy number variation regions with a read depth of < 0.5 or > 1.5 called from *CNVnator* v0.3.3 (Abyzov et al. 2011) were filtered out.

The *AlleleDB* pipeline was run on 864 ChIP-seq datasets in the 6 cell lines from the ENCODE project. In addition to the standard steps in *AlleleDB*, our ASB set was refined by performing beta-binomial tests only within reads overlapping their corresponding TF binding peaks called from the same ChIP-seq dataset. In total, 7,530 ASB SNVs were identified from 638 ChIP-seq datasets.

The ASB SNVs were treated as positive examples in our random forest model. To generate a comparable negative set, we included SNVs from three sources: 1. The 55,611 non-allelic TF binding SNVs, defined by having equal ChIP-seq read counts on two alleles at heterozygous site. 2. The closest variants from each of the SNVs in positive set and outside ChIP-seq peaks (6,373 unique variants in total). 3. A randomly selected set of 1000 genome variants scored no hits on functional annotations from RegulomeDB v1.1. Those three negative sets were combined and weighted equally in our model.

### Building random forest models

For TURF generic scores, seven binary and eight numeric features were created for each variant in the training set (Supplemental Table S1). The seven binary features represent if the variant position overlaps corresponding functional genomic regions by querying RegulomeDB 2.0. Custom scripts were written to retrieve annotations from the RegulomeDB web server. The maximum information content change from PWM was calculated based on the query. Quantiles and variations in ChIP-seq signals pre-calculated from all available bigwig files in ENCODE and functional significance scores from DeepSEA were also incorporated. A random forest model was trained to make predictions on the probability of a query variant being functional. The *scikit-learn 0*.*20*.*3* python package was used to train the random forest model, setting the number of trees to 500.

For TURF tissue-specific scores, a separate random forest model was built with 7 binary tissue-specific features (see feature list in Supplemental Table S1). When training with each ASB cell line, the ASB SNVs in the corresponding cell line were labeled as positive variants, while the other variants were labeled as controls. The *scikit-learn 0*.*20*.*3* python package was used, setting the class_weight option as ‘balanced’.

### Generic scores performance assessment

We evaluated our generic model performance on an independent dataset from an MPRA assay in GM12878 (Tewhey et al. 2018). The labels of the MPRA variants (435 positive variants, 2670 control variants) and prediction scores from DeepSEA (Zhou and Troyanskaya 2015) and regBase were downloaded from regBase database (Zhang et al. 2019). The performance of different tools was assessed on the Area Under ROC Curve (AUROC) and the Area Under Precision-Recall Curve (AUPR).

### Tissue-specific scores performance assessment

The tissue-specific model’s performance was evaluated first on three MPRA datasets in GM12878 (E116), HepG2 (E118) and K562 (E123). The labels for the MPRA variants were obtained from GenoNet (He et al. 2018). The authors labeled the MPRA variants in GM12878 from (Tewhey et al. 2018) with a slightly different criteria than regBase (Zhang et al. 2019), resulting in 293 positive variants and 2772 control variants. The MPRA variants in HepG2 and K562 were from (Kheradpour et al. 2013), where 524 positive variants and 1439 control variants were in HepG2, and 339 positive variants and 1361 control variants were in K562. The same evaluation process as described in GenoNet (He et al. 2018) was used to compare TURF to other available tools, including DeepSEA (Zhou and Troyanskaya 2015), CADD (Kircher et al. 2014) and GenoSkyline (Lu et al. 2016). In detail, we calculated AUROC, AUPR and the correlation coefficient using 1000 replicates of 4:1 random partition of each MPRA dataset. For the divided five parts, four parts were used for training while the remaining part was used for testing.

When evaluating performance with allele specific TF binding SNVs, pre-calculated scores from GenoNet (He et al. 2018) and GenoSkyline (Lu et al. 2016) were downloaded from https://zenodo.org/record/3336209/files/ and http://zhaocenter.org/GenoSkyline.

### Extension to organ-specific scores

The mapping from tissues and cell types (i.e. biosamples) to organ names was downloaded from the ENCODE website (https://www.encodeproject.org/report/?type=BiosampleType). When generating organ-specific prediction scores, we combined the annotations from functional genomics data in all biosamples belonging to the corresponding organ. 51/55 organs had available ChIP-seq data of histone marks and DNase-seq data to generate organ-specific scores.

### Organ-specific significance scores

We calculated organ-specific significance scores relative to a background set from GWAS variants. The GWAS variants were downloaded and assigned to their mapped traits from the GWAS Catalog (Buniello et al. 2019). SNVs on chromosomes 1-22 and chromosome X were the only ones considered for the organ-specific scoring. Linkage disequilibrium (LD) expansion was performed by including SNVs from the 1000 genome project that are in strong LD (R^2^ threshold of 0.6, precalculated R^2^ values downloaded from gs://genomics-public-data/linkage-disequilibrium) with any GWAS SNV. To convert each organ-specific score to a significance score, we calculated the portion of GWAS variants with a greater score in the corresponding organ and did a negative log10 transformation on to the portion (Fig. 5).

### Organ-specific scores enrichment of GWAS traits

In the enrichment analysis, we focused on the GWAS traits with the enrichment of regulatory variants, which have at least 20 GWAS SNVs and at least 5% of the LD-expanded GWAS SNVs in the trait that have TURF generic scores no less than 0.8 (400 traits in total).

To test the enrichment of organ-specific regulatory variants, each GWAS trait set was first sampled with an equal sized background set from all GWAS SNVs from any trait. Subsequent LD expansion was performed on both the trait set and background set (with a more strict R^2^ threshold of 0.8). To reduce the dependencies across SNVs within each set, the SNVs were pruned on each organ individually so that no two SNPs were within 1MB of each other in the same set. Each SNV in decreasing order on organ-specific score was considered, and only retained a SNV if there was no other SNV within 1Mb. After the pruning process, a p-value was computed from the Mann–Whitney U test for each organ-trait combination, with the alternative hypothesis as SNVs in the trait set have greater organ-specific scores than the background set. This test was repeated by sampling 100 versions of the background set and a total of 100 p-values were obtained for each organ-trait pair. 159 traits had at least one organ passing multiple test correction with an FDR of 5%, applied with the Holm-Sidak test from the python package *statsmodels* v0.12.1.

To determine the top organ for each trait, overall high scores of the trait were compared to other organs. The negative log-transformed p-values from the U tests were used to compute the z-score of each organ over all 51 organs. The mean z-scores over 100 iterations for each organ-trait pair were calculated and hierarchical clustering on the 51 organs was performed using the ward linkage method. The final heatmap (Fig. 6 and Supplemental Fig. S5) only shows organ-trait pairs with a z-scores mean higher than 0 and passing multiple test correction (FDR threshold of 5%).

**Figure 6.**
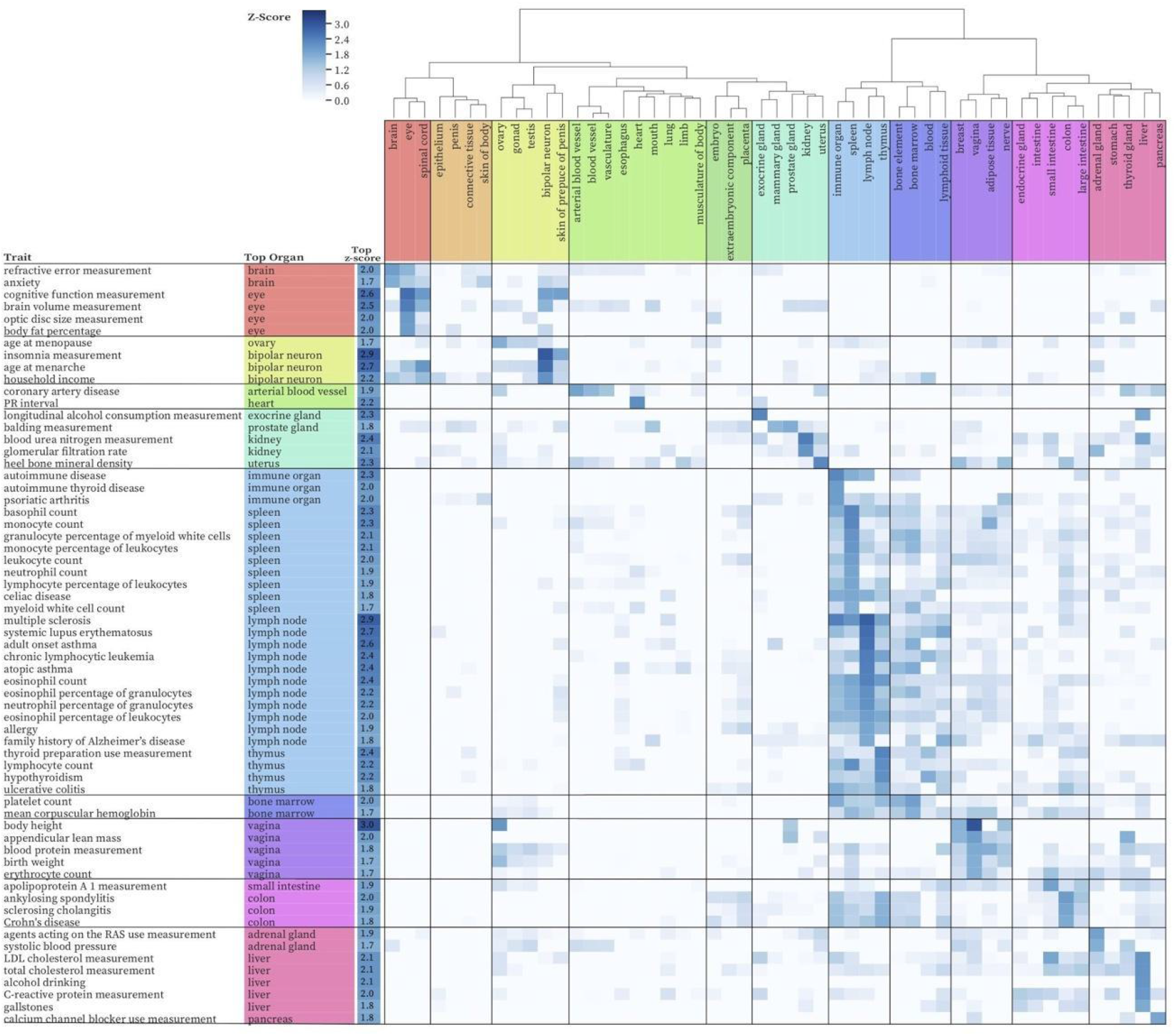
Enrichment of regulatory variants with high organ-specific scores over variants associated with diverse traits. The z-scores of organs (column) for a given trait (row) are shown. The organ with the highest z-score for each trait is shown in additional columns on left. Only organ-trait pairs with z-scores higher than 0 and passing multiple test correction (FDR threshold of 5%) are shown. Traits with z-scores < 1.7 were ignored in this plot, see complete plot Supplemental Figure S5.

## Data access

Pre-calculated TURF generic and organ-specific scores on GWAS variants and data used in this study are available at: https://github.com/Boyle-Lab/RegulomeDB-TURF.

## Acknowledgments

This project was supported by NIH U24 HG009293 to A.P.B. We thank all members in the Boyle lab for helpful discussions and constructive feedback. We thank Yunhai Luo and Ben Hitz for help in retrieving annotations from the RegulomeDB web server.

## Notes

### Competing Interest Statement

The authors have declared no competing interest.

## References

1000 Genomes Project Consortium, Auton A, Brooks LD, Durbin RM, Garrison EP,Kang HM, Korbel JO, Marchini JL, McCarthy S, McVean GA, et al. 2015. A global reference for human genetic variation. Nature 526: 68–74.

Abyzov A, Urban AE, Snyder M, Gerstein M. 2011. CNVnator: an approach to discover,genotype, and characterize typical and atypical CNVs from family and population genome sequencing. Genome Res 21: 974–984.

Alipanahi B, Delong A, Weirauch MT, Frey BJ. 2015. Predicting the sequence specificities of DNA- and RNA-binding proteins by deep learning. Nat Biotechnol 33: 831–838.

Backenroth D, He Z, Kiryluk K, Boeva V, Pethukova L, Khurana E, Christiano A,Buxbaum JD, Ionita-Laza I. 2018. FUN-LDA: A Latent Dirichlet Allocation Model for Predicting Tissue-Specific Functional Effects of Noncoding Variation: Methods and Applications. Am J Hum Genet 102: 920–942.

Barski A, Cuddapah S, Cui K, Roh T-Y, Schones DE, Wang Z, Wei G, Chepelev I, Zhao K. 2007. High-resolution profiling of histone methylations in the human genome. Cell 129: 823–837.

Bernstein BE, Stamatoyannopoulos JA, Costello JF, Ren B, Milosavljevic A, Meissner A,Kellis M, Marra MA, Beaudet AL, Ecker JR, et al. 2010. The NIH RoadmapEpigenomics Mapping Consortium. Nat Biotechnol 28: 1045–1048.

Boyle AP, Davis S, Shulha HP, Meltzer P, Margulies EH, Weng Z, Furey TS, Crawford GE. 2008. High-resolution mapping and characterization of open chromatin across the genome. Cell 132: 311–322.

Boyle AP, Hong EL, Hariharan M, Cheng Y, Schaub MA, Kasowski M, Karczewski KJ,Park J, Hitz BC, Weng S, et al. 2012. Annotation of functional variation in personalgenomes using RegulomeDB. Genome Res 22: 1790–1797.

Buniello A, MacArthur JAL, Cerezo M, Harris LW, Hayhurst J, Malangone C, McMahon A, Morales J, Mountjoy E, Sollis E, et al. 2019. The NHGRI-EBI GWAS Catalog of published genome-wide association studies, targeted arrays and summarystatistics 2019. Nucleic Acids Res 47: D1005–D1012.

Chen J, Rozowsky J, Galeev TR, Harmanci A, Kitchen R, Bedford J, Abyzov A, Kong Y,Regan L, Gerstein M. 2016. A uniform survey of allele-specific binding and expression over 1000-Genomes-Project individuals. Nat Commun 7: 11101.

Davis CA, Hitz BC, Sloan CA, Chan ET, Davidson JM, Gabdank I, Hilton JA, Jain K,Baymuradov UK, Narayanan AK, et al. 2018. The Encyclopedia of DNA elements(ENCODE): data portal update. Nucleic Acids Res 46: D794–D801.

Degner JF, Pai AA, Pique-Regi R, Veyrieras J-B, Gaffney DJ, Pickrell JK, De Leon S,Michelini K, Lewellen N, Crawford GE, et al. 2012. DNase I sensitivity QTLs are amajor determinant of human expression variation. Nature 482: 390–394.

DePristo MA, Banks E, Poplin R, Garimella KV, Maguire JR, Hartl C, Philippakis AA, del Angel G, Rivas MA, Hanna M, et al. 2011. A framework for variation discovery and genotyping using next-generation DNA sequencing data. Nat Genet 43: 491–498.

Diehn JJ, Diehn M, Marmor MF, Brown PO. 2005. Differential gene expression in anatomical compartments of the human eye. Genome Biol 6: R74.

Dong S, Boyle AP. 2019. Predicting functional variants in enhancer and promoter elements using RegulomeDB. Hum Mutat 40: 1292–1298.

ENCODE Project Consortium. 2012. An integrated encyclopedia of DNA elements inthe human genome. Nature 489: 57–74.

Glickman RM, Green PH. 1977. The intestine as a source of apolipoprotein A1. Proc Natl Acad Sci U S A 74: 2569–2573.

He Z, Liu L, Wang K, Ionita-Laza I. 2018. A semi-supervised approach for predicting cell-type specific functional consequences of non-coding variation using MPRAs. Nat Commun 9: 5199.

Johnson DS, Mortazavi A, Myers RM, Wold B. 2007. Genome-wide mapping of in vivo protein-DNA interactions. Science 316: 1497–1502.

Kasowski M, Grubert F, Heffelfinger C, Hariharan M, Asabere A, Waszak SM, Habegger L, Rozowsky J, Shi M, Urban AE, et al. 2010. Variation in transcription factor binding among humans. Science.

Kasowski M, Kyriazopoulou-Panagiotopoulou S, Grubert F, Zaugg JB, Kundaje A, Liu Y,Boyle AP, Zhang QC, Zakharia F, Spacek DV, et al. 2013. Extensive variation in chromatin states across humans. Science.

Kelley DR, Reshef YA, Bileschi M, Belanger D, McLean CY, Snoek J. 2018. Sequential regulatory activity prediction across chromosomes with convolutional neuralnetworks. Genome Res 28: 739–750.

Kheradpour P, Ernst J, Melnikov A, Rogov P, Wang L, Zhang X, Alston J, Mikkelsen TS,Kellis M. 2013. Systematic dissection of regulatory motifs in 2000 predicted human enhancers using a massively parallel reporter assay. Genome Res 23: 800–811.

Kircher M, Witten DM, Jain P, O’Roak BJ, Cooper GM, Shendure J. 2014. A general framework for estimating the relative pathogenicity of human genetic variants. Nat Genet 46: 310–315.

Li MJ, Li M, Liu Z, Yan B, Pan Z, Huang D, Liang Q, Ying D, Xu F, Yao H, et al. 2017. cepip: context-dependent epigenomic weighting for prioritization of regulatory variants and disease-associated genes. Genome Biol 18: 52.

Li MJ, Wang LY, Xia Z, Sham PC, Wang J. 2013. GWAS3D: Detecting human regulatory variants by integrative analysis of genome-wide associations, chromosome interactions and histone modifications. Nucleic Acids Res 41: W150–8.

Lu Q, Powles RL, Wang Q, He BJ, Zhao H. 2016. Integrative Tissue-Specific Functional Annotations in the Human Genome Provide Novel Insights on Many Complex Traits and Improve Signal Prioritization in Genome Wide Association Studies. PLoS Genet 12: e1005947.

Maurano MT, Humbert R, Rynes E, Thurman RE, Haugen E, Wang H, Reynolds AP,Sandstrom R, Qu H, Brody J, et al. 2012. Systematic localization of common disease-associated variation in regulatory DNA. Science 337: 1190–1195.

Mikkelsen TS, Ku M, Jaffe DB, Issac B, Lieberman E, Giannoukos G, Alvarez P,Brockman W, Kim T-K, Koche RP, et al. 2007. Genome-wide maps of chromatin state in pluripotent and lineage-committed cells. Nature 448: 553–560.

Musunuru K, Strong A, Frank-Kamenetsky M, Lee NE, Ahfeldt T, Sachs KV, Li X, Li H,Kuperwasser N, Ruda VM, et al. 2010. From noncoding variant to phenotype viaSORT1 at the 1p13 cholesterol locus. Nature 466: 714–719.

Patwardhan RP, Hiatt JB, Witten DM, Kim MJ, Smith RP, May D, Lee C, Andrie JM, Lee S-I, Cooper GM, et al. 2012. Massively parallel functional dissection of mammalian enhancers in vivo. Nat Biotechnol 30: 265–270.

Quang D, Xie X. 2016. DanQ: a hybrid convolutional and recurrent deep neural network for quantifying the function of DNA sequences. Nucleic Acids Res 44: e107.

Roadmap Epigenomics Consortium, Kundaje A, Meuleman W, Ernst J, Bilenky M, Yen A, Heravi-Moussavi A, Kheradpour P, Zhang Z, Wang J, et al. 2015. Integrative analysis of 111 reference human epigenomes. Nature 518: 317–330.

Rozowsky J, Abyzov A, Wang J, Alves P, Raha D, Harmanci A, Leng J, Bjornson R,Kong Y, Kitabayashi N, et al. 2011. AlleleSeq: analysis of allele-specific expression and binding in a network framework. Mol Syst Biol 7: 522.

Sherry ST. 2001. dbSNP: the NCBI database of genetic variation. Nucleic Acids Research 29: 308–311.http://dx.doi.org/10.1093/nar/29.1.308.

Shigaki D, Adato O, Adhikari AN, Dong S, Hawkins-Hooker A, Inoue F, Juven-Gershon T, Kenlay H, Martin B, Patra A, et al. 2019. Integration of multiple epigenomic marks improves prediction of variant impact in saturation mutagenesis reporter assay. Hum Mutat 40: 1280–1291.

Song L, Crawford GE. 2010. DNase-seq: a high-resolution technique for mapping active gene regulatory elements across the genome from mammalian cells. Cold Spring Harb Protoc 2010: db.prot5384.

Tam V, Patel N, Turcotte M, Bossé Y, Paré G, Meyre D. 2019. Benefits and limitations of genome-wide association studies. Nat Rev Genet 20: 467–484.

Tewhey R, Kotliar D, Park DS, Liu B, Winnicki S, Reilly SK, Andersen KG, Mikkelsen TS, Lander ES, Schaffner SF, et al. 2018. Direct Identification of Hundreds of Expression-Modulating Variants using a Multiplexed Reporter Assay. Cell 172:1132–1134.

Wang Z, White DL, Hoogeveen R, Chen L, Whitsel EA, Richardson PA, Virani SS,Garcia JM, El-Serag HB, Jiao L. 2018. Anti-Hypertensive Medication Use, Soluble Receptor for Glycation End Products and Risk of Pancreatic Cancer in the Women’s Health Initiative Study. J Clin Med Res 7.http://dx.doi.org/10.3390/jcm7080197.

Ward LD, Kellis M. 2016. HaploReg v4: systematic mining of putative causal variants,cell types, regulators and target genes for human complex traits and disease. Nucleic Acids Res 44: D877–81.

Yang B, Zhou W, Jiao J, Nielsen JB, Mathis MR, Heydarpour M, Lettre G, Folkersen L,Prakash S, Schurmann C, et al. 2017. Protein-altering and regulatory genetic variants near GATA4 implicated in bicuspid aortic valve. Nat Commun 8: 15481.

Zhang S, He Y, Liu H, Zhai H, Huang D, Yi X, Dong X, Wang Z, Zhao K, Zhou Y, et al.2019. regBase: whole genome base-wise aggregation and functional prediction for human non-coding regulatory variants. Nucleic Acids Res 47: e134.

Zhou J, Troyanskaya OG. 2015. Predicting effects of noncoding variants with deeplearning-based sequence model. Nat Methods 12: 931–934.

